# A mechanistic model of the S-shaped population growth

**DOI:** 10.1101/012104

**Authors:** Lev V. Kalmykov

## Abstract

The main idea of this note is to show the most basic and purely mechanistic model of population growth, which has been used by us to create models of interspecific competition for verification of the competitive exclusion principle (*1*, *2*). Our logical deterministic individual-based cellular automata model demonstrates a spatio-temporal mechanism of the S-shaped population growth.

---

A classical model of the S-shaped population growth is the Verhulst model. Unfortunately, this is completely non-mechanistic (black-box) model as the internal structure of the complex system and mechanisms remain hidden (*3*). Here I show a completely mechanistic ‘white-box’ model of the S-shaped population growth (Fig. 1 and Movie S1).

**Figure 1.**
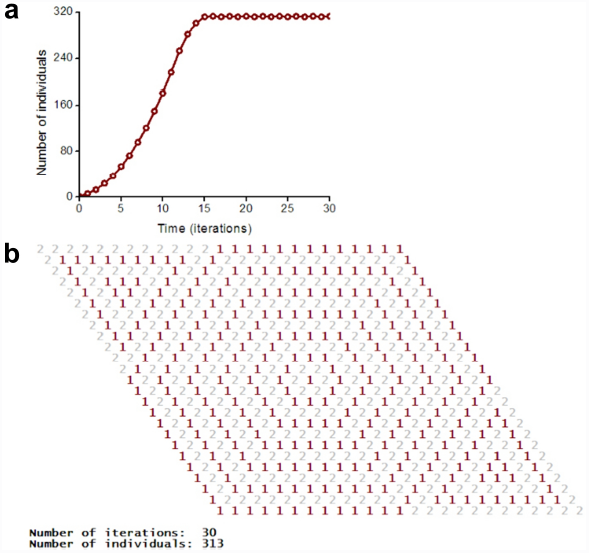
S-shaped population growth. A logical deterministic individual-based cellular automata model of single species population dynamics (Movie S1). **a**, S-shaped population growth curve. **b**, Cellular automata lattice at the 30^th^ iteration.

**A biological prototype of the model** is aggressive vegetative propagation of rhizomatous lawn grasses – e.g. Festuca rubra trichophylla (Slender creeping red fescue). One individual corresponds to one tiller (Fig. 2). A tiller is a minimal semi-autonomous grass shoot that sprouts from the base. Rhizomes are horizontal creeping underground shoots using which plants vegetatively (asexually) propagate. Unlike a root, rhizomes have buds and scaly leaves. One tiller may have maximum six rhizoms in the model. A tiller with roots and leaves develops from a bud on the end of the rhizome. A populated microhabitat goes into the regeneration state after an individual’s death. The regeneration state of the site corresponds to the regeneration of microhabitat’s resources including recycling of a dead individual (Fig. 2b). All individuals are identical. Propagation of offsprings of one individual leads to colonization of the uniform, homogeneous and limited habitat. Finite size of the habitat and intraspecific competition are the limiting factors of the population growth. The maximum possible number of offsprings of one individual is six (Fig. 2a). An individual may propagate in all nearest microhabitats according to the logical rules (Figs 2 and 3).

**Figure 2.**
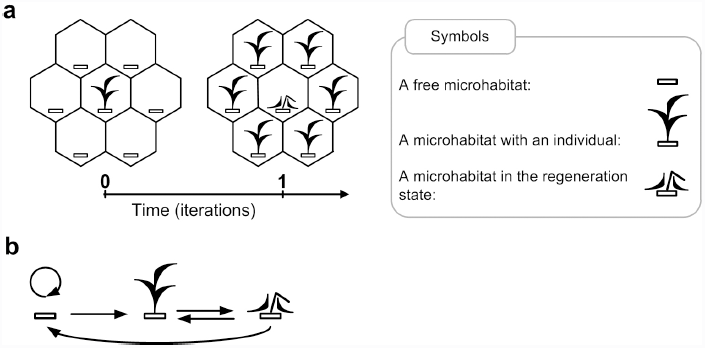
Biological interpretations of the model. **a**, Identical offsprings of the one parental individual occupy all nearest microhabitats what corresponds to aggressive vegetative propagation of plants. The maximum number of offsprings per one individual equals six. The neighbourhood defines fecundity and spatial positioning of offsprings. **b**, A biological interpretation of the graph of transitions between the states of a lattice site. The graph represents a birth-death-regeneration process.

**A mathematical description of the model**. A cellular automata model is defined by the 5-tuple:

1. a lattice of sites;
2. a set of possible states of a lattice site;
3. a neighborhood;
4. rules of transitions between the states of a lattice site;
5. an initial pattern.

Rules of the cellular automata model are presented in Fig. 3 and in the following text.

**Figure 3.**
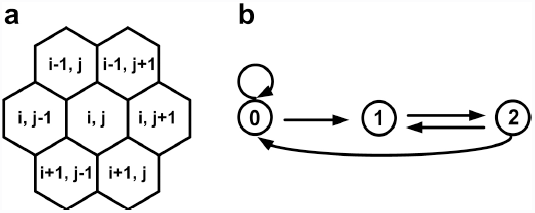
Rules of the cellular automata model. **a**, Hexagonal neighborhood. Coordinates i and j are integer numbers. **b**, Directed graph of transitions between the states of a lattice site.

The lattice consists of 25x25 sites and it is closed on the torus to avoid boundary effects (Fig. 1b). Each site may be in one of the three states 0, 1 or 2, where:

**0** – a free microhabitat which can be occupied by an individual of the species;
**1** – a microhabitat is occupied by a living individual of the species;
**2** – a regeneration state of a microhabitat after death of an individual of the species.

A free microhabitat is the intrinsic part of environmental resources per one individual and it contains all necessary resources and conditions for an individual’s life. A microhabitat is modeled by a lattice site. The cause-effects relations are logical rules of transitions between the states of a lattice site (Fig. 3):

**0→0**, a microhabitat remains free if there is no one living individual in its neighborhood;
**0→1**, a microhabitat will be occupied by an individual of the species if there is at least one individual in its neighborhood;
**1→2**, after death of an individual of the species its microhabitat goes into the regeneration state;
**2→0**, after the regeneration state a microhabitat becomes free if there is no one living individual in its neighborhood;
**2→1**, after the regeneration state a microhabitat is occupied by an individual of the species if there is at least one individual in its neighborhood.

**Physically speaking** this is the simplest model of active (excitable) media with autowaves (travelling waves, self-sustaining waves) (*1*, *4*, *5*). An active medium is a medium that contains distributed resources for maintenance of autowave. An autowave is a self-organizing dissipative structure. An active medium may be capable to regenerate its properties after local dissipation of resources. In our model, reproduction of individuals occurs in the form of population waves (Fig. 1). We use the axiomatic formalism of Wiener and Rosenblueth for simulation of excitation propagation in active media (*6*). In accordance with this formalism rest, excitation and refractoriness are the three successive states of a site. In our model the rest state corresponds to the free state of a microhabitat, the excitation state corresponds to the life activity of an individual in a microhabitat and the refractory state corresponds to the regeneration state of a microhabitat. All states have identical duration. If the refractory period will be much longer than the active period, then such a model may be interpreted, for example, as propagation of the single wave of fire on the dry grass. Time duration of the basic states can be easily varied using additional states of the lattice sites.

According to Alexander Watt, a plant community may be considered ‘*as a working mechanism*’ which ‘*maintains and regenerates itself*’ (*7*). This logical model of the single-species population dynamics shows such mechanism in the direct and most simplified form. We consider the white-box modeling by logical deterministic cellular automata as a perspective way for investigation not only of population dynamics but also of all complex systems (*1*, *3*). The main feature of this approach is the use of cellular automata as a way of linking semantics (ontology) and logic of the subject area. Apparently, the effectiveness of this approach is provided by the fact that cellular automata are an ideal model of time and space.

## Acknowledgements

I thank Vyacheslav L. Kalmykov for useful discussions and suggestions.

